# A Novel Pan-Proteome Array for High-Throughput Profiling of the Humoral Response to *Treponema pallidum* subsp. *pallidum*: a Pre-Clinical Study

**DOI:** 10.1101/2024.04.20.590429

**Authors:** Joseph J. Campo, Emily R. Romeis, Amit Oberai, Jozelyn V. Pablo, Christopher Hung, Andy A. Teng, Adam D. Shandling, Amber Phan, Austin M. Haynes, Lorenzo Giacani

## Abstract

**Background:** Given the resurgence of syphilis, research endeavors to improve current assays for serological diagnosis and management of this disease are a priority. A proteome-scale platform for high-throughput profiling of the humoral response to *Treponema pallidum* (*T. pallidum*) proteins during infection could identify antigens suitable to ameliorate the performance and capabilities of treponemal tests (TTs), which may require weeks to become positive following infection, cannot distinguish between active and previously treated infections, or assess treatment response. Additionally, because infection-induced immunity is partially protective, profiling the response to *T. pallidum* outer membrane proteins (OMPs) could help select vaccine candidates.

**Methods:** We developed a pan-proteome array (PPA) based on the Nichols and SS14 strain complete proteomes and used it to define the IgM and IgG humoral response to 1,009 *T. pallidum* proteins in sera collected longitudinally from long-term infected rabbits, and from rabbits that were infected, treated, and re-infected.

**Findings:** Approximately a third of the pathogen’s proteome was recognized in infected animals, with a marked IgG response detectable between day-10 and day-20 post-infection. We found early, gradual, and late IgG kinetic profiles, strain-dependent differences in humoral reactivity, and post-treatment fluctuation in reactivity for several antigens. Very few antigens elicited an IgM response. Several OMPs were significantly and differentially recognized, but few elicited a robust response.

**Interpretation:** The PPA allowed the identification of antigens that could facilitate early diagnosis and of a core set of OMP that could explain protection upon re-infection. No antigen appeared suitable to monitor treatment response.

**Funding:** NIH SBIR-R43AI149804

**RESEARCH IN CONTEXT:** *Evidence before this study:* In April 2024, we searched the PubMed database for articles on preclinical studies using high throughput proteome arrays containing at least 10% of the predicted *T. pallidum* proteome that aimed at identifying antibody reactivity to *T. pallidum* antigens during experimental syphilis infection. We could retrieve only one manuscript. In this work, an array containing the *T. pallidum* partial proteome as annotated in the first sequenced Nichols strain genome (GCA_000008605.1) in 1998 was assembled using recombinant antigens expressed in *Escherichia coli* (*E. coli*). The resulting array was probed using pooled sera from three rabbits infected with the Nichols stain of *T. pallidum*, attained from infected animals at five time points following intratesticular infection. The small number of reactive antigens (n = 106) identified in this early study was likely to be an incomplete set of all antigens recognized during infection because not all the predicted targets in the *T. pallidum* proteome were successfully expressed and tested. In retrospect, additional limitations of the study included an initial suboptimal annotation of the Nichols genome used to define the pathogen’s proteome, which has now changed with the availability of a re-sequenced Nichols strain genome devoid of sequencing errors that affected the initial annotation process, and the refinement of bioinformatic pipelines for the identification of open reading frames (ORFs). Furthermore (as acknowledged by the authors), the possible presence of amplification errors in their expression clones might have affected the sequence of some protein targets and antibody binding to the targets. As a result, some of the *T. pallidum* antigens known to elicit a robust humoral response during experimental infection were not detected in this antigenic screen. Lastly, employing only the Nichols strain in this early study did not consider that a significant portion of the circulating syphilis strains belong to the SS14 clade of *T. pallidum*.

*Added value of this study:* This novel PPA, combined with a more robust experiential design than ever reported, allowed us to overcome most of the limitations associated with the study mentioned above, as we were able to a) use the most recent annotations for the selected *T. pallidum* strains based on accurate genome sequences, b) print the pathogen’s virtually complete proteome in the study array, c) analyze individual sera to account for rabbit-to-rabbit variability in the humoral response to infection rather than pooled sera, d) detect both IgM and IgG over 10 or 20 timepoints, depending on the experimental design, e) obtain information on how the humoral response evolved upon treatment and re-infection and, finally, f) evaluate all of the above in animals infected with two *T. pallidum* strains whose genetic background is representative of the two currently circulating clades of the syphilis agent.

*Implications of all the available evidence:* Our study provides new and more comprehensive data on how humoral immunity for two classes of antibodies develops during infection and how it evolves in response to treatment and re-infection. The analysis of sera collected at tightly spaced time points post-inoculation and for an extensive period post-infection provides a wealth of information to improve the diagnostic performance of existing tests detecting treponemal antigens. The analysis of differential immunity specific to the pathogen’s putative OMPs provides a rationale for vaccine candidate selection.

## INTRODUCTION

Syphilis is a multistage chronic sexually transmitted infection that is still a significant burden for public health despite being treatable. The World Health Organization (WHO) estimated the syphilis global prevalence and incidence to be between 18-36 million cases and 5.6-11 million new annual cases, respectively ^1,2^. Although most cases occur in low- and middle-income countries, syphilis rates have also steadily increased for two decades in high-income countries ^3–8^. In the US, the rate of primary and secondary syphilis in 2022 (17.7 cases per 100,000 population) represented a 743% increase compared to the 2.1 cases per 100,000 population reported in 2000 ^3^. Similarly, the rate of total (all stages) syphilis increased from 11.2 (in 2000) to 62.2 cases per 100,000 population in 2022 ^3^. If untreated, syphilis might progress to affect the cardiovascular and central nervous systems, potentially leading to severe manifestations such as aortic aneurism, stroke-like syndrome, dementia, and paralysis ^9^. Because the syphilis agent, *Treponema pallidum* subsp. *pallidum* (*T. pallidum*), can cross the placenta, mother-to-child transmission of the infection during pregnancy can lead to stillbirth, perinatal death, and a plethora of other adverse pregnancy outcomes, with severe repercussions for maternal and infant health. In sub-Saharan Africa, for example, ∼50% of stillbirths are predicted to be related to congenital syphilis ^10^. Even in the US, where maternal health standards are very high, syphilis resurgence in women of reproductive age has led to 3,755 cases of congenital syphilis reported by the CDC in 2022 ^3^, which is a substantial increase compared to the 554 cases on record in 2000. Syphilis global epidemiology strongly supports the need for novel research endeavors to curtail the spread of this serious infection. In addition to identifying alternative treatments to benzathine penicillin G (BPG) ^11^ and developing an effective vaccine for syphilis ^12^, research efforts should aim at improving diagnostic approaches to both reduce the temporal gap between exposure and diagnosis to minimize the consequences of *T. pallidum* dissemination to virtually all bodily organs from the site of infection, and assess treatment efficacy.

Laboratory diagnosis of syphilis is mainly achieved through serology, with a testing algorithm encompassing the use of two separate and complementary tests ^13^. Lipoidal serologic tests, such as the Venereal Disease Research Laboratory (VDRL), detect antibodies to lipoidal molecules (mainly cardiolipin, lecithin, and cholesterol) released by both damaged host cells and *T. pallidum* during active infection ^14^, and they are generally referred to as non-treponemal tests (NTT, albeit this is now considered a misnomer). Seroconversion, however, occurs 3-6 weeks after exposure to the pathogen. The relatively low sensitivity of these assays (74-99%) sometimes causes failure to detect infection in patients with primary syphilis. The fluctuation of serum titer of lipoidal tests is, however, important to monitor successful response to treatment, relapse, or re-infection following a syphilis episode. An effective antibiotic therapy should induce a fourfold decrease in lipoidal test titer within 3-12 months (depending on the infection stage at diagnosis), which is indicative of serological cure. Despite proper treatment, however, serological non-responders might occur among patients with late syphilis, and lipoidal tests might remain reactive at a low level for years (a state known as serofast) ^15^. If a history of syphilis is known, a rise in lipoidal test titer generally supports re-infection or treatment failure and the need for additional treatment.

A positive lipoidal test result is verified with a treponemal test (TT) to detect antibodies to specific *T. pallidum* antigens, such as the *T. pallidum* Particle Agglutination assay (TPPA), the Fluorescent Treponemal Antibody Absorption test (FTA-ABS) or the Chemiluminescence and Enzyme Immunoassays (CIA, EIA). Treponemal tests like the TPPA and FTA-ABS are performed manually and use whole *T. pallidum* as the source of antigen (either intact or sonicated), while automated high-throughput assays like the EIA and CIA employ recombinant *T. pallidum* lipoproteins, known to be highly specific for *T. pallidum* subspecies, abundantly expressed, and immunodominant. Although TTs have excellent sensitivity and specificity, their efficacy is still limited in the diagnosis of early syphilis ^13^, do not discriminate between active or past infections like lipoidal tests do, and cannot be used to assess syphilis stage, even though limited research suggests that differential immune responses to specific *T. pallidum* antigens might occur during infection progression and in response to therapy ^16^.

Developing and testing an antigen array encompassing the whole *T. pallidum* proteome could lead to the identification and recruitment of additional antigens to increase the sensitivity and overall capabilities of TTs based on recombinant proteins, and perhaps provide evidence on whether a TT could allow for disease staging based on differential immunoreactivity over time, or help monitor treatment response, relapse, or re-infection in previously treated patients.

Furthermore, it is established that long-term-infected rabbits (>90 days) develop partially protective immunity which leads to attenuation of disease manifestations upon re-infection ^17,18^, and clinical studies have suggested that patients who experienced a syphilis episode are more likely to be asymptomatic with a subsequent episode ^19–24^. These findings strongly suggest that an effective syphilis vaccine may be attainable if the right antigens are targeted. Because evidence exists that antibodies to *T. pallidum* surface-exposed outer membrane proteins (OMPs) are significant mediators of protective immunity ^25–27^, the identification of the OMP that preferentially elicit a humoral response during experimental infection could provide clues regarding candidates to include in an experimental vaccine design.

In this study, we constructed a pan-proteomic array (PPA) based on the annotated proteomes of the Nichols and SS14 *T. pallidum* isolates, which are the model laboratory strains for the two genetically distinct clades of this pathogen currently circulating worldwide ^28^. The PPA was tested with sera collected longitudinally from long-term Nichols- and SS14-infected animals and from animals that were infected with either Nichols or SS14, treated, and then re-infected with the homologous strain two months post-treatment. Here, we report on the portion of *T. pallidum* proteome recognized during experimental infection with either strain, on the IgG and IgM antibody profiles that develop in infected rabbits, on the antibody kinetic profiles in long-term infected animals and in rabbits that were re-infected following effective treatment with BPG.

## METHODS

### Ethics statement and animal care

New Zealand White (NZW) rabbits were used for treponemal propagation and experimental infections to obtain serum samples over time. Adult male NZW rabbits were purchased from Western Oregon Rabbit Company (Philomath, OR) and housed at the Animal Use and Care Facility (ARCF) at the UW. After a period of acclimation, blood was collected from the central ear artery to perform both a lipoidal and treponemal serological test and rule out past or current infection with *Treponema paraluiscuniculi*, the known agent of rabbit syphilis. More specifically, heat-inactivated sera from these rabbits were tested individually using the VDRL test (BD, Franklin Lanes, NJ) test and the SERODIA TPPA assay (Fujirebio Diagnostics, Inc., Malvern, PA) according to the provided protocols. Only VDRL- and TPPA-negative rabbits were enrolled in this study. Animal care was provided in accordance with the Guide for the Care and Use of Laboratory Animals, and all experimental procedures were conducted under protocol #4243-01 (PI: Lorenzo Giacani), approved by the University of Washington (UW) IACUC. All animal procedures were conducted in compliance with the ARRIVE guidelines. Rabbit euthanasia was performed consistent with the recommendations of the Panel on Euthanasia of the American Veterinary Association.

### Strain Propagation and DNA extraction

The *T. pallidum* Nichols and SS14 strains were propagated through intratesticular (IT) inoculation in NZW rabbits as previously described ^29^ to obtain treponemal cells as source of DNA to construct the PPA and, subsequently, to produce the inoculum for all experimental infections. Briefly, 10^7^ treponemes were injected in each testis. Once peak orchitis was reached (about 10 days post-IT infection with Nichols treponemes and 20-25 days for the SS14 strain), animals were euthanized, and testes were removed and minced in 10 mL of sterile saline. The resulting suspension was spun for 10 min at 1,000 x g in a 5430 Eppendorf (Hauppauge, NY) centrifuge at room temperature to remove cellular debris. Treponemes in the supernatant were enumerated using dark-field microscopy (DFM) and diluted to the appropriate concentration to inoculate naïve rabbits IT for further propagation or intradermally (ID) for the actual study. Treponemes harvested for DNA extraction were spun for 10 min at 15,000 rpm at 4°C using an Eppendorf 5425 refrigerated centrifuge. Pellets were resuspended in 400 µl 1X Lysis Buffer (10mM Tris pH 8.0, 0.1M EDTA pH 8.0, 0.5% SDS) and frozen at −80°C until extraction using the QIAamp DNA Mini Kit (Qiagen Inc., Chatsworth, CA) according to the provided protocol.

### Experimental infection, treatment, and re-infection

For the experimental infections, 24 rabbits divided into two groups of twelve animals were used.

One group was infected with the Nichols strain, and the other with the SS14 strain. Each animal was infected intradermally (ID) on six sites on their shaved backs with 10^6^ treponemes per site. Each group was then divided into sub-groups of six animals. The first subgroup of Nichols-infected animals was labeled Nichols long-term (N.LT) because these animals were infected and followed longitudinally for 90 days, during which serum samples were collected at day 5 and day 10 post-infection and, subsequently, every ten days until day 90 post-infection, when the animals were bled from the central ear artery one last time and then euthanized. Pre-infection sera were obtained at the time the animals arrived at the vivarium. The second subgroup of six Nichols-infected animals was labeled Nichols re-infection (N.RI) because animals were treated at day 30 post-inoculation with 200,000 units of BPG (Bicillin L-A, Pfizer, New York, NY) administered intramuscularly (IM) in a single injection, and equivalent w/w to 4.8 million units for humans. Two months post-treatment (at study day 90), the animals were re-infected as described above with the homologous Nichols strain. Re-infected animals were followed longitudinally for three additional months. From these animals, serum samples were obtained before initial inoculation, at day 5 and day 10 post-initial infection and every ten days after for the remainder of the 180-day experiment. On day 90 after re-infection (at study day 180), the animals were bled and euthanized. The same experimental design described for the N.LT and N.RI rabbits was applied to the 12 animals infected with the SS14 strains, which were identified as the SS14 long-term (S.LT; n = 6 animals), and SS14 re-infection (S.RI; n = 6 animals) groups, respectively. All 24 rabbits enrolled in this study were shaved regularly in order not to hinder dermal lesion development. Collected sera were heat-inactivated at 56°C for 30 min and aliquoted. One aliquot was used for syphilis serology, while the remaining ones were frozen at −80°C to be used later to probe the PPA.

### VDRL and TPPA Serology

The VDRL and TPPA tests were used post-inoculation with *T. pallidum* to assess the establishment and progression of the infection. Both tests were performed according to the provided protocols. When sera tested positive by VDRL, serum titration was also performed. In addition, TPPA-reactive sera collected at day 30 and day 90 (from both LT and RI animals), and at day-180 (RI animals only, as the LT rabbits were euthanized at day 90 post-infection) were also titrated.

The technologist was blinded to the treatment status and group of the animals from which the samples were collected. Mean log2 titers ± SE were calculated for each group at each time point: nonreactive = −1.0; weakly reactive = −0.5; reactive at a titer of 1:1= 0; reactive at a titer of 1:2 = 1, and so on. Antilogs of mean ± SE of titers were plotted with nonreactive set at y = 0. Following testing, serum samples were stored at −80°C until use. Throughout the study, only three serum samples could not be collected, all from the N.LT group (from two rabbits at day 80 post-inoculation and from one rabbit at day 90 post-infection). Statistical analysis was performed using Student’s t-test with significance set at *p*<0.05.

### Array construction and probing

A core proteome was identified using the genome sequences of the Nichols (NC_021490.2/CP004010.2) and SS14 (NC_021508.1/CP004011.1) strains. Pan-proteome arrays were fabricated containing 1,040 full-length or fragmented recombinant proteins representing a total of 1,009 genes. Of these, 986 were genes encompassing the core proteome shared by the two strains, 10 were genes encoding for Nichols-specific proteins and 13 were specific for the SS14 strain. Each open reading frame (ORF) sequence was amplified by PCR, inserted into the vector pXT7 by recombination in *E. coli* to establish a library of partial or complete coding DNA sequences. Proteins were expressed using a coupled *E. coli* cell-free *in vitro* transcription and translation (IVTT) system (Rapid Translation System, Biotechrabbit, Berlin, Germany) and spotted onto nitrocellulose-coated glass AVID slides (Grace Bio-Labs Inc., Bend, OR) using an Omni Grid Accent robotic microarray printer (Digilabs Inc. Hopkinton, MA). Each expressed protein included a 5’ poly-histidine (His) epitope and 3’ hemagglutinin (HA) epitope. Pan-proteome array chip printing and protein expression were quality checked by probing random slides with anti-His and anti-HA monoclonal antibodies fluorescently labeled and quantifying spot signals using a microarray scanner. As positive control antigens, six recombinant *T. pallidum* proteins previously expressed to construct a minimal proteomic array ^30^ or other experimental purposes were spotted onto the array at two different concentrations. Those controls included the known immunodominant lipoproteins Tp0435/TpN17, Tp0574/TpN47, and Tp0768/TmpA, known to be conserved in both strains, the putative OMPs Tp0117/TprC and Tp0131/TprD2, and a relatively conserved amino-terminal fragment of the Tp0897/TprK antigen, also known to be targeted by the host response during infection based on previous studies ^31^. Except for TprK, all proteins were printed on the array as full-length molecules based on their available annotation.

Serum samples were diluted (1:100) and incubated on the PPA chips overnight at 4°C on a rocker. Bound IgG was detected with DyLight650-conjugated goat anti-rabbit IgG (Bethyl Laboratories, Montgomery, TX), and IgM was detected with DyLight550-conjugated goat anti-rabbit IgM (Abcam Inc., Cambridge, UK). Washed and dried PPA chips were scanned, and the spot and background signal intensities (SI) were exported into R package for statistical analysis. Sera collected from the LT and the RI animal groups were probed in separate batches.

### Statistical Analysis

Spot SIs were adjusted for local background by subtraction, and values were floored to 1. Next, the data were normalized by dividing the protein spot values by the median of IVTT control spots (IVTT expression reactions with no *T. pallidum* ORFs), and values were log transformed using the base-2 logarithm. Thus, normalized data represented the log2 signal-to-noise ratio, where a value of 0 represents specific antibody SI equal to the background, 1.0 represents twice the background, 2.0 represents 4-fold over background, and so forth. *T. pallidum* protein responses were classified as seropositive for SI of at least 1.0, or twice the background. Maximal SIs across all timepoints for each rabbit were calculated for each protein. A protein was classified as “reactive” if over half of the rabbits responded to the protein, i.e., median max SI > 1. Individual antibody responses or mean antibody responses by study day were visualized using the ComplexHeatmap package ^32^. The distributions of reactive antigen responses were visualized using error bar plots of all proteins. Overlap in reactivity to individual antigens between groups was visualized using the UpSetR package, which recasts the information of a complex Venn diagram as bar plots ^33^. The association of protein physiochemical and functional features with antibody reactivity was analyzed using logistic regression within the safeBinaryRegression package ^34^ with separation = ‘find’ to identify terms separating the observations when maximum likelihood estimate is found not to exist, followed by performing stepwise regression on the previous model using the stepAIC function set to both forward and backward stepwise regression. Only protein features present in at least 5 antigens and did not return singularity errors were included. Differences in antibody levels between sequential time points were assessed using paired t-tests. Analysis was stratified by Nichols and SS14 strain infection groups, and the LT and RI experiments were analyzed separately. Differences between time points were visualized using volcano plots, and longitudinal trajectories of specific antibody responses were visualized using line plots with the ggplot2 package ^35^. Only reactive antigens were included in differential analysis, and p-values were adjusted for the false discovery rate using the method described by Benjamini and Hochberg ^36^.

## RESULTS

### VDRL and TPPA Serology

Over time, all rabbits in both the long-term-infected N.LT and S.LT groups became VDRL positive, even though rabbits inoculated with the Nichols strain seroconverted earlier than those inoculated with the SS14 strain (Fig.1A). More specifically, all the N.LT rabbits were VDRL-positive by day 20 post-inoculation, while all S.LT rabbits were positive by day 40 post-inoculation (Fig.1A). The highest VDRL titer was 1:16 (for three and four animals in the N.LT and S.LT groups, respectively). By day 50 post-inoculation, titers had declined in most N.LT animals, while a decline began at later time points for the S.LT rabbits. These results are consistent with an active infection established in all animals. Consistently, all the rabbits in the N.RI group (Fig.1B) were VDRL-positive by day 20 post-inoculation, whereas only 2 (33%) rabbits in the S.RI group (Fig.1B) seroconverted before treatment (day 30). All of the N.RI and S.RI VDRL-positive rabbits seroreverted to a nonreactive VDRL within a month following treatment administration. Upon animal re-infection (60 days post-BPG treatment; experiment day 90), an increase in the mean VDRL reactivity was measurable starting at day 140 for the animals in the N.RI group (∼50 days after re-inoculation; Fig.1B), and day 160 for the SS14-infected animals (∼70 days after re-inoculation; Fig.1B). At the end of the experiment (day 180 post-infection), all N.RI and S.RI animals had seroconverted, albeit titers remained lower in S.RI rabbits than in rabbits from the N.RI group. These results support that infection was established, cured, and established again in the RI animal groups.

**Figure 1.**
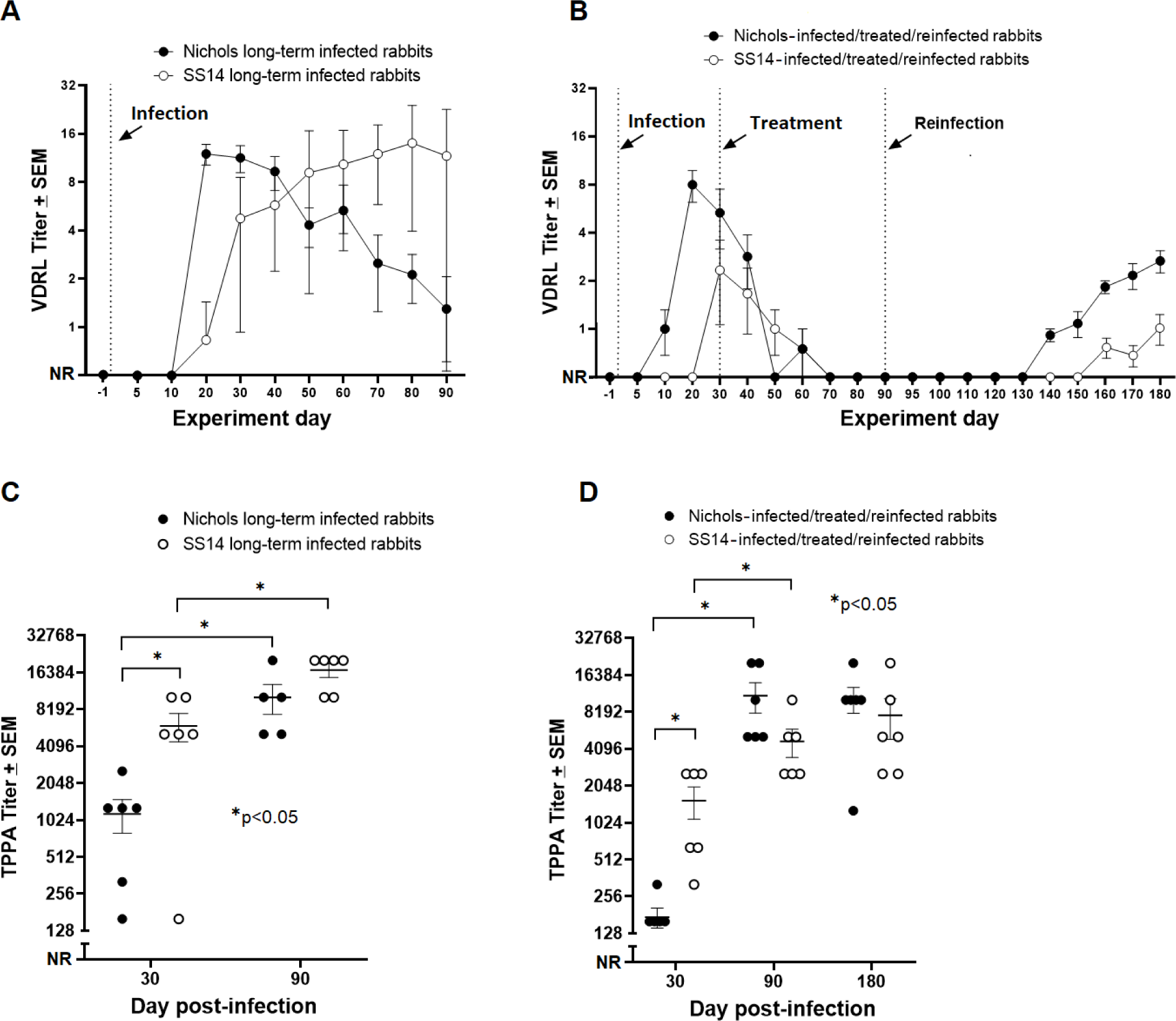
Serological test results for rabbits enrolled in this study. **(A)** VDRL titers for long-term (LT) infected rabbits inoculated with either the Nichols or SS14 strain measured over 90 days post-infection. Vertical dotted line indicates the infection event (experiment day 0) **(B)** VDRL titers for rabbits in the reinfection (RI) groups inoculated with either the Nichols or SS14 strain and measured over 180 days post-infection. Vertical dotted lines mark the infection, treatment, and re-infection events (experiment day 0, 30, and 90, respectively). For calculating the mean ± SE, titers were converted to log2 with nonreactive = −1.0; weakly reactive = −0.5; reactive 1:1 = 0, reactive 1:2 = 1, and so forth. Antilogs of mean ± SE of titers are shown in this figure with nonreactive set at y = 0. **(C)** TPPA titers for long-term (90 days) infected rabbits inoculated with either the Nichols or SS14 strain measured at day 30 and day 90 post-infection. **(D)** TPPA titers for rabbits in the RI groups inoculated with either the Nichols or SS14 strain measured at day 30, 90 (time of treatment), and 180 post-infection.

All infected rabbits in the N.LT and S.LT groups were TPPA-positive at day 30 post-inoculation and displayed significantly higher mean titers at day 90 post-infection than at day 30 (Fig.1C), which was consistent with infection progression. The mean titer was significantly higher in the S.LT group than in the N.LT group at day 30, but not significantly different between the two groups at day 90 post-infection. All the N.RI and S.RI rabbits had seroconverted by day 30 post-inoculation (Fig.1D), supporting establishment of the infection also in the three S.RI animals that did not become VDRL-positive before treatment (Fig.1B). Compared to day 30, TPPA titers were significantly higher also for the N.RI and S.RI groups at the time of re-infection (day 90), but not significantly different between the two groups at day 90 post-infection. Day 180 titers were not significantly different from day 90 titers for both groups (Fig.1D). Overall, these serological data further support that inoculation with *T. pallidum* led to a productive infection in all animals.

### Profiling of the humoral responses in long-term infected rabbits

Overall, sera collected from the N.LT rabbits showed that a total of 207 antigens over 1008 unique targets (encompassing core and Nichols-specific proteins) were significantly recognized over the course of the infection (Fig.2A). Positive targets represented approximately ∼20% of the Nichols strain proteome (as annotated in NC_021490.2). At the end of the 3-month infection period, sera from S.LT animals recognized a total of 327 antigens, corresponding to ∼31% of the SS14 proteome (as annotated in NC_021508.1; Fig.2B). In N.LT animals, there were no significant increases in specific IgG responses to *T. pallidum* antigens by 10 days post-infection, although there was a non-significant positive trend for a few antigens (Fig.2C). By day 30, there were 79 antibody responses that significantly increased from day 10 (Fig.2D). From day 30 to day 60, no additional antibody responses were significantly increased based on our set false discovery rate (FDR), except for Tp0409a/YajC preprotein translocase subunit. At this time point, however, a non-significant positive trend was detectable for numerous antigens (Fig.2E), represented by circles above the dotted line). The change in the number of targeted proteins observed between day 60 and day 90 post-infection was not significant (Fig.2F). Animals infected long-term with the SS14 strain (S.LT) also had no significant changes by day 10 post-infection (Fig.2G), while 94 antibody responses were significantly increased from day 10 to day 30 (Fig.2H), and 3 from day 30 to day 60 (Fig.2I), corresponding to the hypothetical proteins Tp0565 and Tp0895, and the Tp0499/MreD rod-shape determining protein. As for the N.LT animals, no significant changes in antibody levels were observed between day 60 and day 90 post-infection in S.LT animals (Fig.2J), aside from a positive trend for a few targets. The discrepancy between total number of reactive antigens and number of significantly increased antibodies between timepoints is explained by the statistical power to detect incremental increases in antibody levels between sequential 10-day intervals and variation in the numbers of reactive antigens at different time points, as opposed to reactivity by maximal responses across the study period.

**Figure 2.**
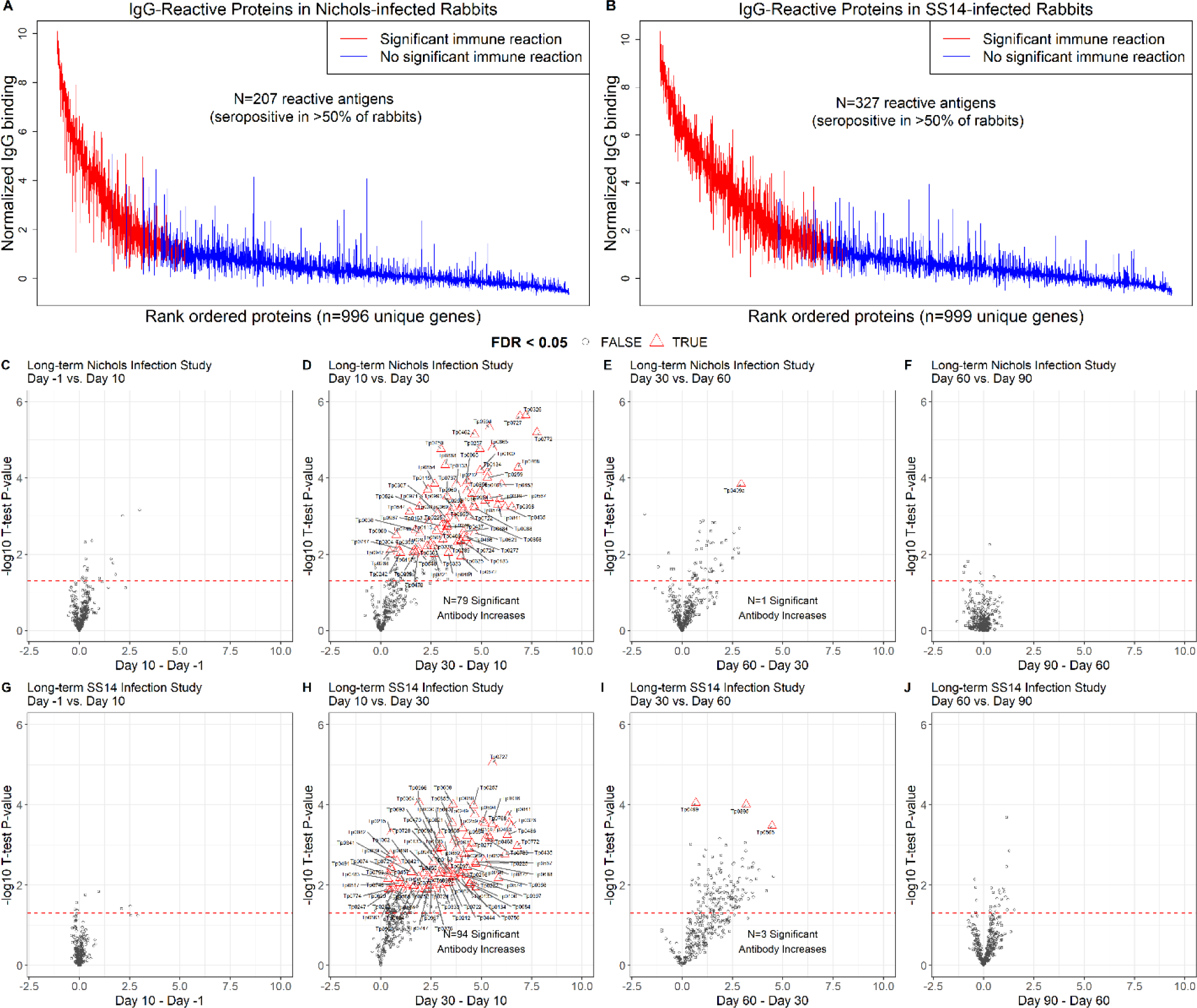
IgG reactivity and longitudinal trajectory in long-term (N.LT and S.LT) infected rabbits. (**A-B**) Interquartile range plots showing the maximal normalized IgG binding signal for each rabbit for each *T. pallidum* protein over the 90-day period of the long-term infection. Each bar represents a protein on the array, and proteins are ordered by the median signal of (**A**) Nichols-infected (N.LT) rabbits and (**B**) SS14-infected (S.LT) rabbits. Red bars represent a significantly reactive protein or proteins with seropositive responses in at least four out of six total rabbits in the group. (**C-J**) Volcano plots showing the difference in normalized IgG binding between timepoints for each reactive protein on the array (x-axis) by the inverse log_10_ P-value of paired Student’s T-tests. In each plot, the horizontal red dashed line represents an unadjusted P-value of 0.05. All values above the red line are below 0.05. After adjustment for the false discovery rate, antibody responses that remained statistically significant were highlighted as red triangles and labeled with the antigen annotated identifier. (**C-F)** Plots showing sequential timepoint differences for N.LT animals. **(G-J)** Plots showing sequential timepoint differences for S.LT animals.

The increase in the number of proteins significantly targeted by IgG antibodies began between day 10 and day 20 for both N.LT and S.LT animals and continued through day 30, but were not significantly increased thereafter (Fig.S1), likely due to limits in statistical power. In contrast, the numbers of reactive antigens at each time point (i.e., antigens seropositive in over half of animals at each study day), were similar for N.LT and S.LT animals until day 30, thereafter plateauing for N.LT but continuing to increase until day 60 post-infection for S.LT animals, to then reach plateau (Fig.3). The breadth of reactive antigens with increasing antibody levels was more consistent with the number of reactive antigens when timepoints were compared against baseline (Fig.S2). The complete list of antigens recognized in both the N.LT and S.LT strains and time to a seropositive response is provided in File S1.

**Figure 3.**
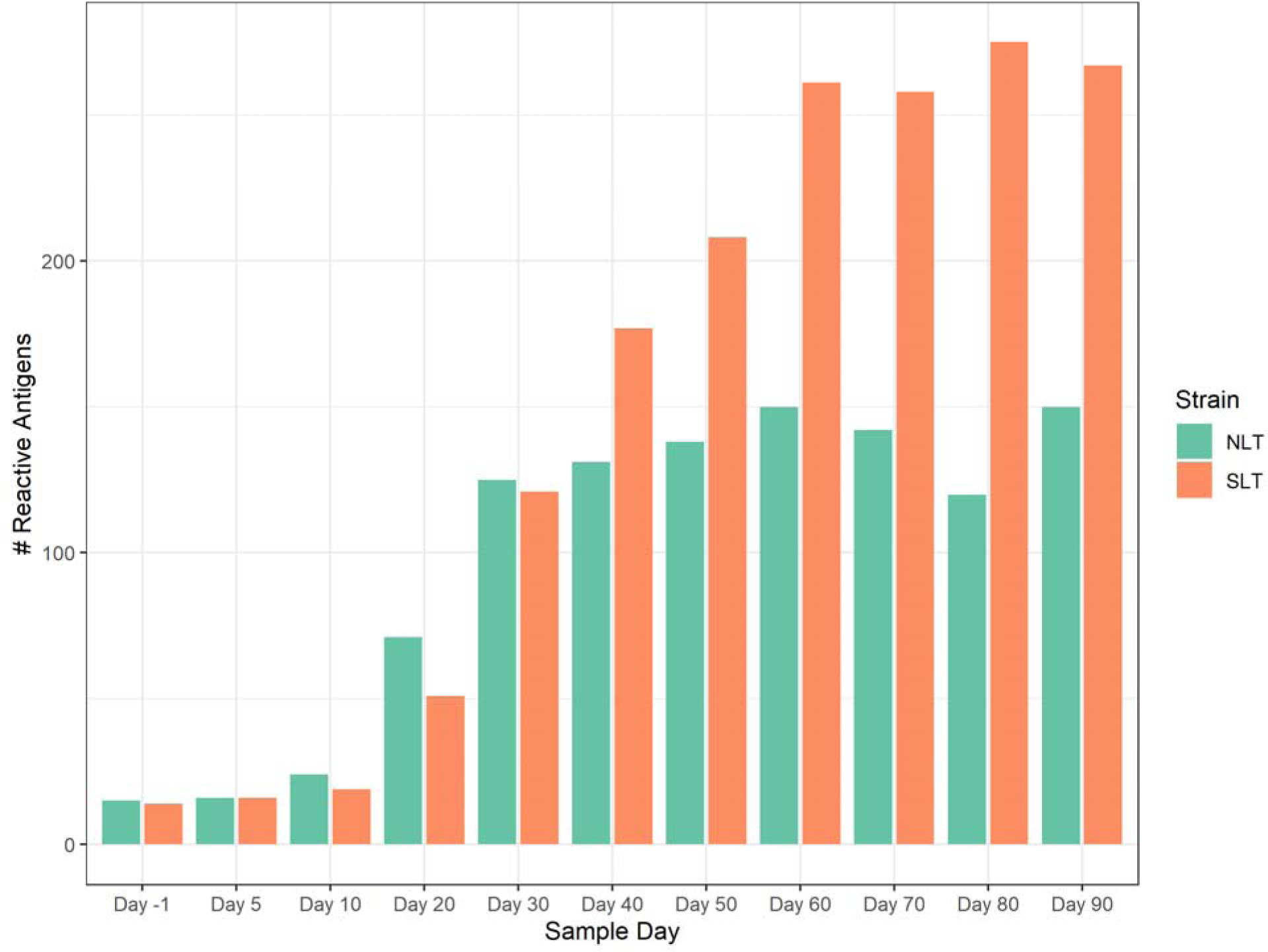
Antigens with significant immune responses at each time point in long-term (N.LT and S.LT) infected rabbits. Bar plot showing the count of *T. pallidum* proteins with significant reactivity, or proteins with seropositive responses in at least four out of six total rabbits in each group at each given time point. The grouped orange bars represent Nichols-infected (N.LT) animals, while green bars represent long-term SS14-infected (S.LT) animals.

The robust IgG response that developed in both LT rabbit groups was accompanied by a weak IgM response. More specifically, in the N.LT and S.LT infected animals, respectively, a total of 13 proteins elicited a weak but significant IgM response (File S1; Fig.S3A). The rabbits in the N.LT group recognized 11 targets, while the S.LT animals only five (Fig.S3A). In N.LT animals, our stepwise regression model associated significant IgM reactivity to the Tp0319-TmpC, Tp0435-TpN17, and Tp0684-MglB lipoproteins, while the remaining targets were either annotated as hypothetical proteins (Tp0707 and Tp0990) or proteins with random functions such as Tp0154 (one of the three *T. pallidum* pseudouridylate synthase), albeit only at two time points post-inoculation, Tp0567 (PrfB peptide chain release factor II), TP0653 (PotB ABC transporter permease), and Tp0727 (FlgE flagellar hook protein). The only putative OMP recognized was Tp0733 (an OmpW homolog) (Fig.S3A). The S.LT rabbits, on the contrary, only recognized the Tp0435-TpN17 and Tp0768-TmpA lipoproteins, the Tp0772 hypothetical protein, Tp0733-OmpW, and the Tp0567-PrfB factor. In both groups, the Tp0435-Tp17 protein elicited the strongest IgM response (Fig.S3A).

Of the 351 total targets in the array found to have elicited a significant IgG response in the long-term infected animals (regardless of the infecting strain), 182 antigens were recognized by both the Nichols- and SS14-infected rabbits, while 140 antigens were recognized only by sera from the SS14-infected animals, and 19 antigens elicited a response exclusively in Nichols-infected rabbits (Fig.4A; File S1). Stepwise regression identified seven main categories of reactive proteins/enzymes (Fig.4B). In order of significance, such categories were: lipoproteins (p=2.0^−09^), metabolism-associated enzymes (p=6.8^−06^), flagellar assembly proteins (p=6.2^−05^), enzymes specific for pyruvate metabolism (p=2.0^−04^), vaccine candidates (i.e., OMPs; p=1.3^−04^), ribosomal proteins (p=2.9^−02^), and chemotaxis-related proteins (p=2.1^−02^). Antibody kinetics shown for individual rabbits for the 30 most reactive antigens in both N.LT and S.LT groups support the early development of reactivity to these targets, and a limited rabbit-to-rabbit variability (Fig.4C; Table 1). The analysis of reactivity limited to annotated/putative *T. pallidum* lipoproteins and putative OMPs/vaccine candidates (Fig.4D) showed that almost the entirety of these targets was recognized by the S.LT rabbit group, although at various degrees of intensity. The Tp0684-MglB, Tp0435-TpN17, Tp0768-TmpA, Tp0319-TmpC, Tp0574-TpN47, Tp0453 (a concealed outer membrane-associated protein) ^37^, Tp0954 (a putative placental adhesin) ^38^, Tp0971-TpD iron transporter, Tp0993-RlpA, and Tp1016 (a putative lipoprotein with peptidyl prolyl cis/trans isomerase function) were the most significantly recognized lipoproteins, while Tp0326-BamA was the most highly recognized OMP (Fig.4D). Aside from Tp0326-BamA, most OMPs currently evaluated as vaccine candidates elicited a significant but overall weak immunity. Compared to the S.LT rabbits, the Nichols-infected animals failed to recognize the Tp0126-OmpW, Tp0313-TprE, Tp0515-LptD, Tp0621-TprJ, Tp0859-FadL, Tp0969-TolC, Tp01031-TprL OMPs, and the Tp0136 putative outer membrane lipoprotein/adhesin ^39^. Tp0011-TprB was recognized in both strains, but the signal intensity was higher in S.LT rabbits than N.LT ones.

**Figure 4.**
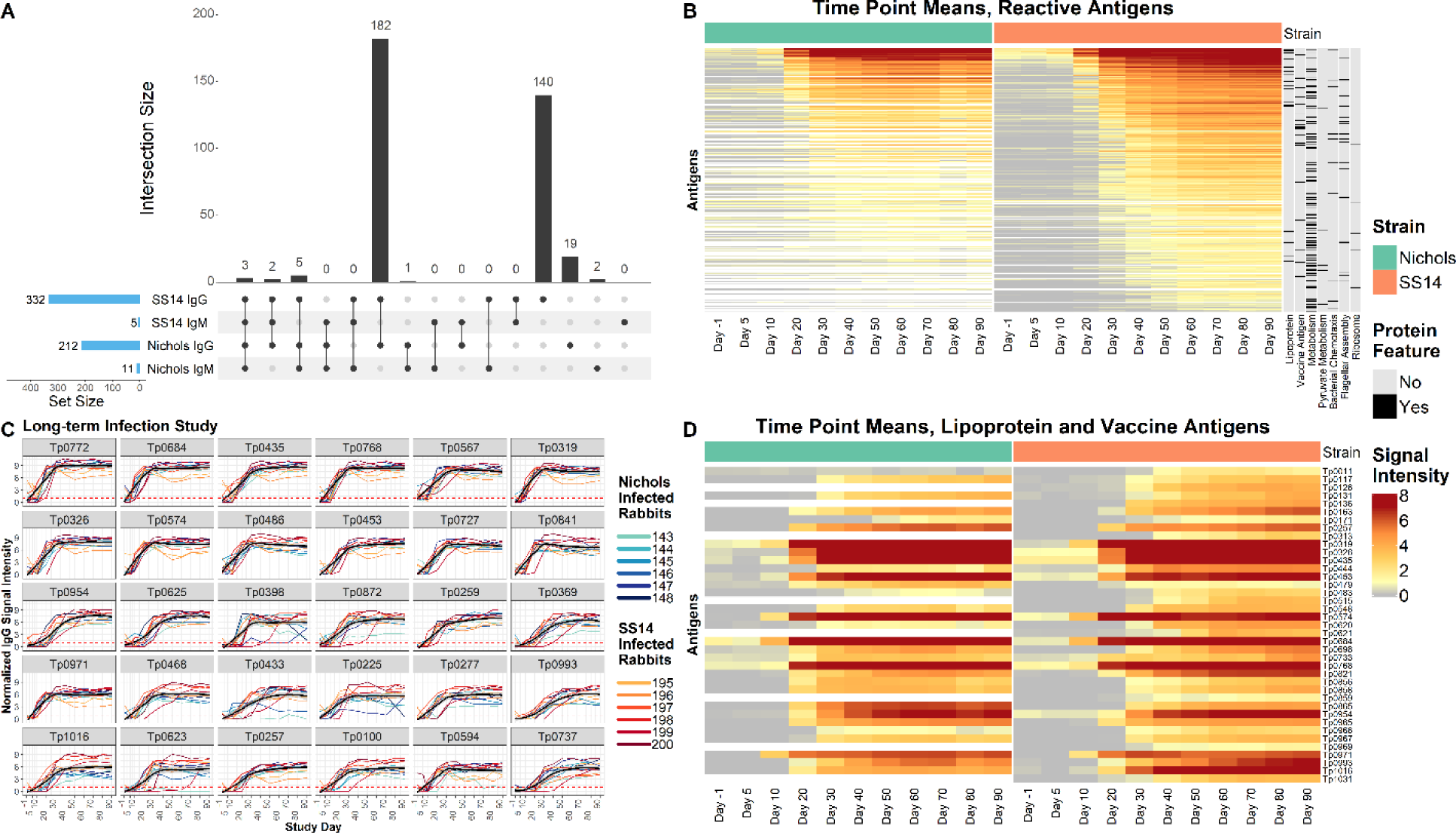
IgG binding profiles in N.LT and S.LT rabbits. **(A)** UpSet plot summarizing both the set and set size of antigens recognized by IgG and IgM during experimental infection of rabbits with either the Nichols or SS14 strains of *T. pallidum*. The UpSet plot is equivalent to a Venn diagram, where each vertical bar represents antibody responses that “intersect” or overlap between two or more categories (i.e., antigens that are reactive in each of the categories specified by the connected dot matrix below the bar graph). The blue horizontal bars represent the total number of reactive antigens in each category. **(B)** Heatmap showing the mean IgG binding Signal Intensity for each reactive antigen in either Nichols or SS14 strain-infected rabbits. Columns represent the means of each group of rabbits at each sequential time point, and rows represent *T. pallidum* proteins sorted by hierarchical clustering. Protein features that were significantly associated with antibody reactivity in stepwise regression models are shown in the black and grey columns to the right of the heatmap. Responses are grouped by Nichols (green header) and SS14 (orange header) strain-infected rabbits. **(C)** Line plots of reactivity to the 30 most reactive antigens in animals infected long-term with the Nichols strain (teal to dark blue lines) and the SS14 strain (orange to dark red lines). In each graph, the black line represents the running mean of all samples at each time point post-infection. **(D)** Heatmap of the antigens annotated as lipoproteins or OMPs/vaccine candidates (ordered by gene ID) recognized in Nichols-infected animals (left side, green header), and SS14-infected rabbits (right side, orange header) over the 90-day infection period.

### Profiling of the humoral responses in treated/re-infected rabbits

A subset of *T. pallidum* antigens elicited a gradual and robust IgG response also in the two groups of RI animals (Fig.5A). More specifically, at the end of the 180-day experiment, following treatment and re-infection, a total of 86 antigens had elicited a significant response in Nichols-infected rabbits, while 99 targets were recognized by rabbits infected with the SS14 strain (Fig.5A). Most of the recognized targets (n = 66) were shared between the two N.RI and S.RI animals, while 29 and 16 proteins, respectively, were recognized only by SS14- or Nichols-infected rabbits (Fig.5A and File S1). Compared to LT animals, fewer targets (n = 4) elicited an IgM response in RI rabbits (Fig.5A and Fig.S3B). More specifically, the N.RI rabbits recognized the Tp0435-TpN17 and Tp0574-TpN47 lipoproteins, as well as the hypothetical protein Tp0707 and the Tp0727-FlgE flagellar hook protein. The S.RI rabbits, on the contrary, only recognized the Tp0707 hypothetical protein (Fig.S3B).

**Figure 5.**
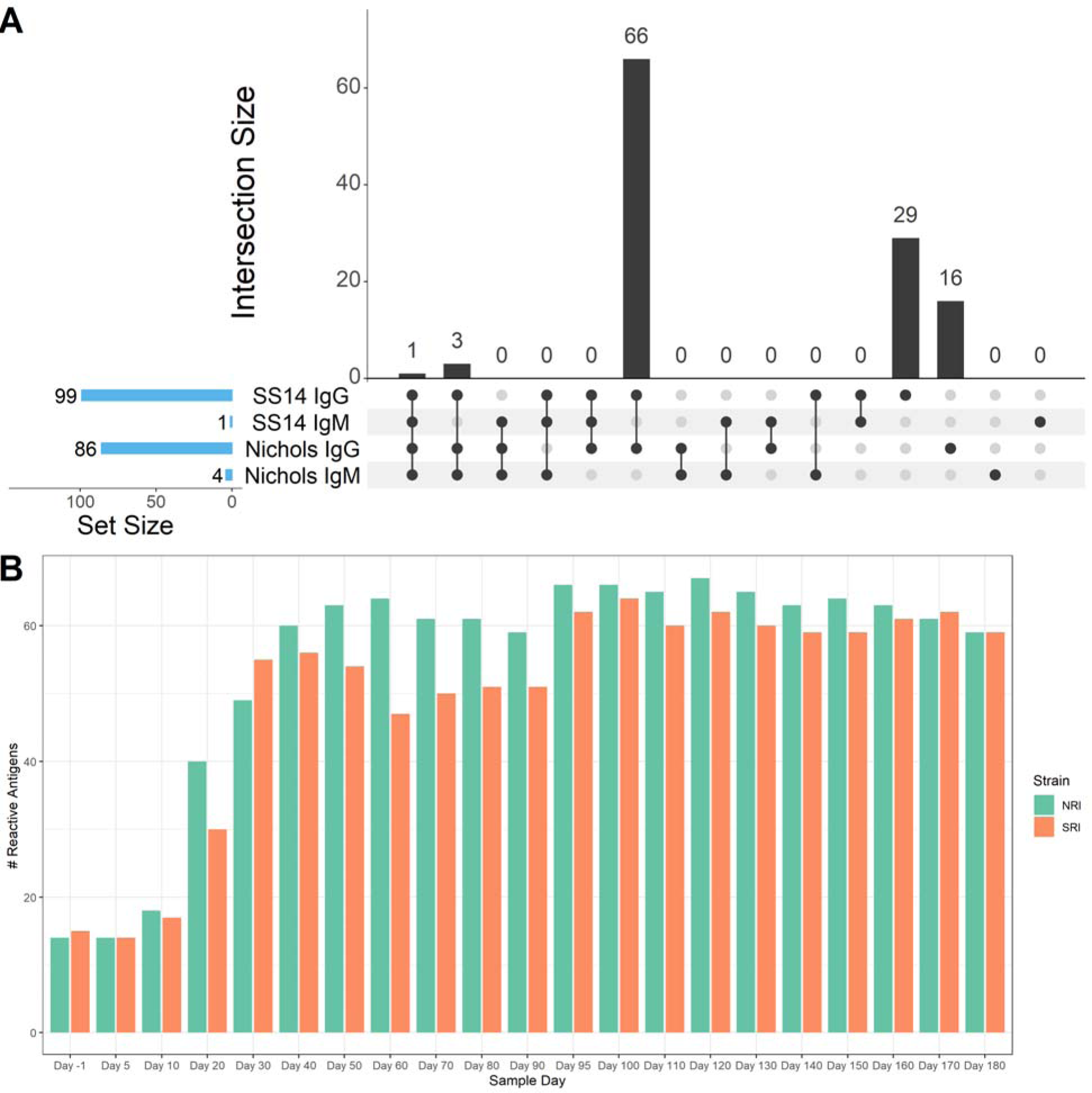
**(A)** Overlap in IgG and IgM antibody responses between N.RI and S.RI rabbit groups. The UpSet plot is equivalent to a Venn diagram, where each vertical bar represents antibody responses that “intersect” or overlap between two or more categories, i.e., antigens that are reactive in each of the categories specified by the connected dot matrix below the bar graph. The blue horizontal bars represent the total number of reactive antigens in each category. **(B)** Antigens with significant immune responses at each time point in the N.RI and S.RI rabbit groups. The bar plot shows the count of *T. pallidum* proteins with significant “reactivity”, or proteins with seropositive responses in at least four of six rabbits at each given time point. The grouped orange bars represent Nichols-infected (N.RI) animals, while green bars represent long-term SS14-infected (S.RI) animals.

**Table 1.** Most significantly recognized *T. pallidum* antigens in S.LT and N.LT rabbits^1^.

| ORF #<sup>2</sup> | Putative gene function | Normalized IgG binding (Nichols-infected rabbits/ SS14-infected rabbits)<sup>3</sup> | Seroprevalence (%) N.LT/S.LT rabbits | Time to 100% seroprevalence (day) N.LT/S.LT rabbits |
| --- | --- | --- | --- | --- |
| Tp0772 | Hypothetical protein/transcriptional regulator | 7.20/7.34 | 100/100 | 20/20 |
| Tp0684 | MglB/glucose-galactose binding lipoprotein | 6.89/7.09 | 100/100 | 10/20 |
| Tp0435 | TpN17 lipoprotein | 6.31/6.90 | 100/100 | 20/20 |
| Tp0768 | TmpA lipoprotein | 6.74/6.91 | 100/100 | 20/20 |
| Tp0567 | FlbB/flagellar protein | 6.37/6.34 | 100/100 | 20/20 |
| Tp0319 | TmpC/membrane lipoprotein; possible ABC transporter solute binding lipoprotein | 6.31/6.47 | 100/100 | 20/10 |
| Tp0326 | BamA/OMP assembly factor | 6.05/6.42 | 100/100 | 20/30 |
| Tp0574 | TpN47 lipoprotein | 6.01/6.03 | 100/100 | 20/10 |
| Tp0486 | Hypothetical protein | 5.54/6.78 | 100/100 | 10/20 |
| Tp0453 | Concealed OMP | 5.69/5.77 | 100/100 | 20/20 |
| Tp0727 | FlgE/flagellar hook protein | 5.68/5.83 | 100/100 | 20/20 |
| Tp0841 | DegQ/serine endoprotease | 5.36/6.06 | 100/100 | 20/20 |
| Tp0954 | Lipoprotein/putative adhesin | 4.78/5.37 | 100/100 | 30/30 |
| Tp0625 | Hypothetical protein | 4.85/5.41 | 100/100 | 30/30 |
| Tp0398 | FliE/flagellar hook-basal body complex protein | 4.29/4.88 | 100/100 | 30/50 |
| Tp0872 | FliD/flagellar filament capping protein | 4.76/5.41 | 100/100 | 30/50 |
| Tp0259 | LysM/peptidoglycan-binding domain-containing protein | 4.99/5.14 | 100/100 | 20/20 |
| Tp0369 | BamD/OMP assembly factor BamD | 3.95/4.65 | 100/100 | 30/40 |
| Tp0971 | TpD/iron transporter lipoprotein | 4.73/5.06 | 100/100 | 10/20 |
| Tp0468 | Tetratricopeptide repeat protein | 4.08/5.36 | 100/100 | 20/30 |
| Tp0433 | Hypothetical protein | 3.77/4.52 | 100/100 | 40/70 |
| Tp0225 | Leucine-rich repeat domain-containing protein | 3.62/5.65 | 100/100 | 20/20 |
| Tp0277 | Ctp peptidase | 3.84/5.25 | 100/100 | 30/30 |
| Tp0993 | RlpA/septal ring lytic transglycosylase | 3.24/4.14 | 100/100 | 30/40 |
| Tp1016 | Peptidyl-prolyl cis-trans isomerase | 2.86/5.40 | 100/100 | 30/30 |
| Tp0618 | Hypothetical protein | 3.31/4.43 | 100/100 | 30/40 |
| Tp0100 | TlpA/disulfide reductase | 3.66/4.71 | 100/100 | 30/40 |
| Tp0257 | GpD/glycerophosphodiester phosphodiesterase | 4.02/4.19 | 100/100 | 20/20 |
| Tp0594 | Hypothetical protein | 3.83/4.29 | 100/100 | 30/40 |
| Tp0737 | Periplasmic binding protein ABC transporter | 3.76/3.46 | 100/100 | 30/50 |

<sup>1</sup> The lineplot is selecting antigens by the means of the maxes for each rabbit. For each protein, the maximum of all timepoints for each rabbit is calculated, and then the mean of those maxes is assigned as the value to sort by.
<sup>2</sup> Based on the annotation of the Nichols (NC\_021490.2/CP004010.2) and SS14 strain (NC\_021508.1/CP004011.1) genomes.
<sup>3</sup> Mean reactivity values over 90 days post-infection.

As seen for the LT rabbits, significant increases in the number of IgG-reactive proteins began between day 10 and day 20 for both N.RI and S.RI animals and continued throughout day 30. Following treatment (day 30 post-infection), the number of reactive antigens increased in N.RI rabbits to reach plateau at experiment day 40, but remained virtually unchanged in the S.RI rabbits. Following successful re-infection (per VDRL serology; Fig.1B), the total number of reactive proteins slightly increased in both animal groups (Fig.5B). The complete list of antigens recognized in both the N.RI and S.RI rabbits and time to a seropositive response is provided in Table S1.

Volcano plots comparing seroreactivity to *T. pallidum* proteins in animals prior to infection and in sera collected at day 30 post-infection (time of treatment) with the Nichols strain showed that significant reactivity had developed to 18 antigens (Fig.6A; Table S1). The same analysis using later timepoint sera supported that the increment in the number of significantly recognized antigens was virtually nonexistent (Fig.6B-D), although a positive trend was seen for numerous proteins (circles above the dotted line). Unlike the animals in the LT groups, rabbits in the S.RI group developed reactivity to a similar number of antigens over the course of the experiment when compared to the N.RI rabbits. Volcano plots comparing the reactivity between different timepoints in the S.RI animals showed that reactivity to most antigens (n = 27) developed before treatment (Fig.6E), and was not significantly increased by re-infection (Fig.6F-H).

**Figure 6.**
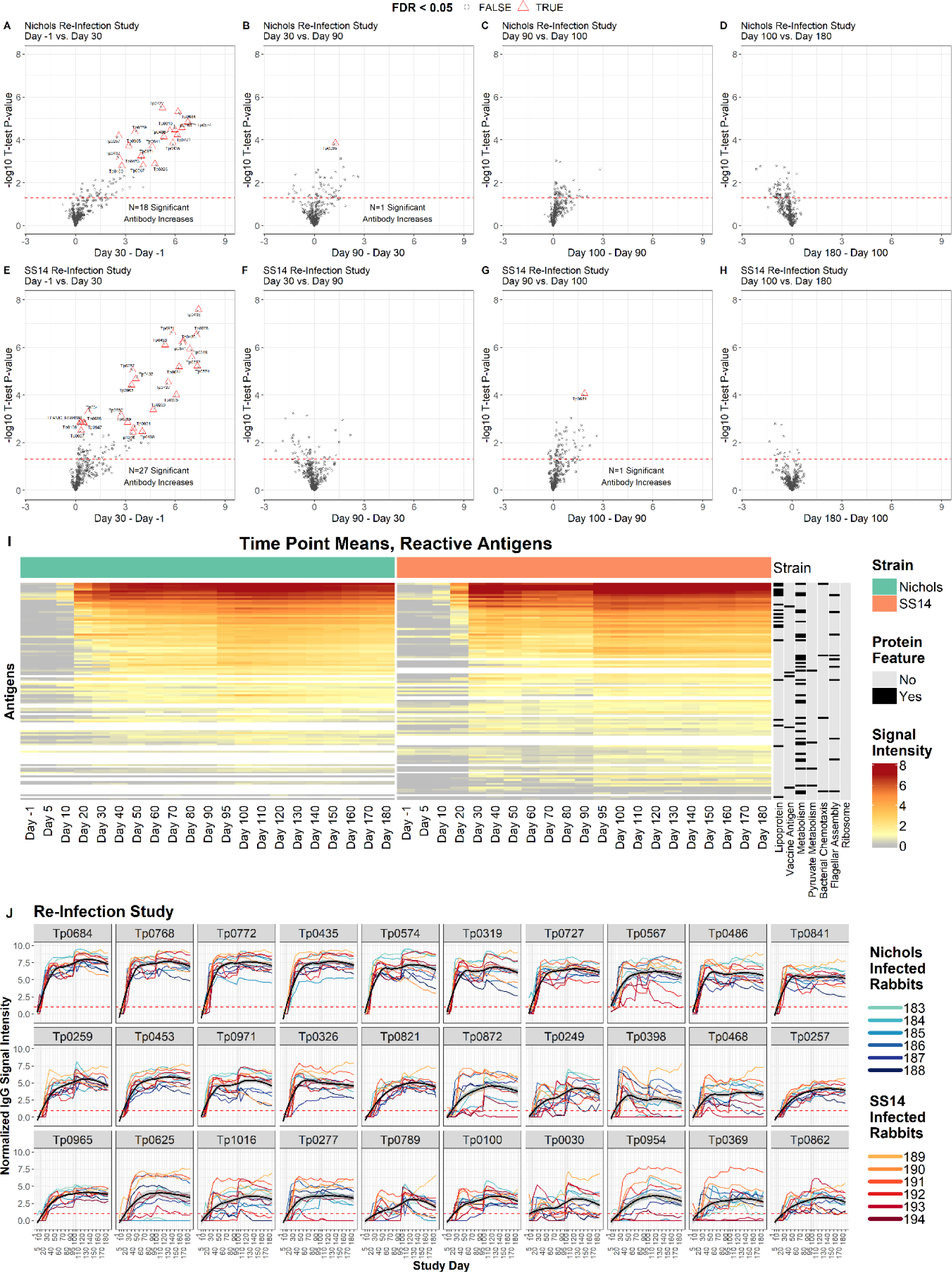
IgG reactivity profiles of rabbits that were infected, treated and then reinfected. **(A-H**) Volcano plots show the difference in normalized IgG binding between timepoints for each reactive protein on the array (x-axis) by the inverse log_10_ P-value of paired Student’s T-tests. The horizontal red dashed lines represent an unadjusted P-value of 0.05, and all values above the red line are below 0.05. After adjustment for the false discovery rate, antibody responses that remained statistically significant were highlighted as red triangles and labeled with the antigen ID. Plots **A-D** are sequential time point differences for Nichols-infected (N.RI) animals, and **E-H** are for SS14-infected (S.RI) animals. **(I)** Heatmap showing the mean IgG binding Signal Intensity for each of the 115 antigens that were reactive in either N.RI or S.RI rabbits. Columns represent the means of each group of rabbits at each sequential time point, and rows represent *T. pallidum* proteins sorted by hierarchical clustering. Protein features that were significantly associated with antibody reactivity in stepwise regression models are shown in the black and grey columns to the right of the heatmap. Responses are grouped by Nichols (green header) and SS14 (orange header) strain-infected rabbits. (**J)** Lineplots showing the longitudinal trajectories of the 30 most reactive antigens by seroprevalence and normalized intensity. Each individual rabbit’s normalized IgG binding signal intensity is plotted as a colored line. Nichols-infected rabbits are teal to dark blue lines, and SS14-infected rabbits are orange to dark red lines. In each graph, the black line represents the running mean of all samples at each time point post-infection. Values are reported in Table S1.

The heatmap of reactive antigens (Fig.6J) graphically shows the effect of BPG treatment (day 30) and re-infection (day 90) in RI animals. Reactivity to several antigens declined over the two months that followed treatment, to then increase again starting at day 95, five days after animal re-inoculation. Overall, a total of 115 proteins were collectively recognized in these two rabbit groups (Fig.5A and File S1). Antibody kinetics shown for individual rabbits for the 30 antigens in both the N.RI and S.RI that were most significantly recognized in both rabbit groups (Fig.6J) showed that a decline in reactivity (black line, representing the running mean of the values form each rabbit from each time point) occurred for several antigens when individual reactivities were averaged, but was never found to be significant at experiment day 90 (at the time of reinfection) compared to experiment day 30, at the time of treatment. Analysis of the kinetic for other antigens, also considering the number of rabbits that seroconverted over the observation period, did not support that any of the reactive targets could be effectively used to monitor treatment response in a fashion comparable to lipoidal serological tests. Table S1 provides reactivity values for each antigen recognized in both the N.RI and S.RI rabbits.

### Antibody response to strain-specific targets

Several targets (Table 2 and File S1) were labeled as strain-specific proteins due to their partial sequence identity between the Nichols and SS14 strains, or because differently annotated by in silico pipelines, like in the case of Tp0040, and Tp0134c. Of the 13 strain-specific targets that elicited a significant IgG response in any of the rabbit groups analyzed in our study (Table 2), there were two putative OMPs, represented by the putative porins Tp0131 (the TprD/TprD2 antigens in Nichols and SS14, respectively), and Tp0548 (a FadL homolog, whose encoding sequence is widely used for *T. pallidum* strain typing) ^40^. Immunity to both Tp0131 variants was detected in both N.LT and S.LT rabbits, although not in the RI animals, likely due to early treatment of the animals and the low-level of expression of OMPs in this pathogen ^41,42^. Consistent with the fact that the TprD/TprD2 variants differ significantly within their central region, that sequence identity exists also with other Tpr paralogs of the syphilis agents (e.g., TprF and TprI) ^43^, and that our array does not have a peptide corresponding uniquely to the TprD2 targeted, however, it is unclear whether the detected immunity is truly specific to Tp0131. The SS14-Tp0548 variant was only recognized in S.LT infected rabbits, while reactivity to the Nichols variant was detected in both N.LT and S.LT animals, supporting that there is enough antigenic similarity for antibody cross-reactivity. The two variants of the Tp0136 putative surface fibronectin binding-protein ^39^ were recognized by both the S.LT and S.RI rabbit groups, but not by any Nichols-infected animals, suggesting that this target might be in general more expressed in the SS14 strain than in Nichols. Overall, none of the SS14- or Nichols-specific antigens exhibit a pattern of recognition fully strain-specific.

**Table 2.** Reactivity to strain-specific antigens.

| Target variant<br>SS14 (S)<br>Nichols (N) | Gene<br>coordinates<sup>1</sup> | Significantly<br>reactive in<br>N.LT vs S.LT | Significantly<br>reactive in<br>N.LT vs S.RI | Annotated<br>function | Percent<br>identity<br>(%) |
| --- | --- | --- | --- | --- | --- |
| Tp0006 (S)<br>Tp0006 (N)<sup>2</sup> | 7014..8261<br>7014..7181 | □□ □□<br>☒ ☒ | □□ ☒<br>☒ ☒ | Hypothetical protein/<br>Probable lipoprotein | 75.4% |
| Tp0027 (S)<br>Tp0027 (N) | 34209..35432<br>34211..35434 | ☒ ☒<br>Not represented<br>in the array<sup>3</sup> | ☒ ☒<br>Not represented<br>in the array<sup>3</sup> | HlyC/CorC family<br>transporter | 99.7% |
| Tp0040 (S)<br>Tp0040 (N) | 46945..49395<br>47597..49390 | □□ ☒<br>□□ ☒ | ☒ ☒<br>□□ ☒ | Methyl-accepting<br>chemotaxis protein | 73.4% |
| Tp0126c (S)<br>Tp0126c (N) | 149381..149866<br>149381..149866 | ☒ ☒<br>☒ ☒ | ☒ ☒<br>☒ ☒ | Hypothetical protein | 88.8% |
| Tp0127 (S)<br>Tp0127 (N) | 149860..150240<br>149809..150477 | ☒ □□<br>□□ □□ | ☒ ☒<br>☒ ☒ | Hypothetical protein | 51.12% |
| Tp0131 (S)<br>Tp0131 (N) | 154110..152314<br>154160..152370 | □□ □□<br>□□ □□ | ☒ ☒<br>☒ ☒ | Putative OMP/porin<br>transporter; TprD/D2 | 79.0 % |
| Tp0134c (S)<sup>4</sup><br>Tp0134c (N) | 156884..157591<br>Not annotated<sup>6</sup> | ☒ □□<br>N/A | ☒ ☒<br>N/A | hypothetical protein | N/A |
| Tp0136 (S)<br>Tp0136 (N) | 157797..157869<br>157747..157819 | ☒ □□<br>☒ □□ | ☒ □□<br>☒ □□ | Putative surface<br>lipoprotein/adhesin | 91.9% |
| Tp0318 (S)<br>Tp0318 (N) | 335991..335812<br>Not annotated<sup>6</sup> | ☒ ☒<br>N/A | ☒ ☒<br>N/A | Hypothetical protein | N/A |
| Tp0461 (S)<br>Tp0461 (N) | 492090..492506<br>492057..492416 | ☒ ☒<br>☒ ☒ | ☒ ☒<br>☒ ☒ | Helix-turn-helix<br>transcriptional regulator | 81.5% |
| Tp0470 (S)<br>Tp0470 (N) | 499709..498768<br>499845..498736 | □□ □□<br>□□ □□ | ☒ ☒<br>□□ □□ | TPR domain protein | 84.8% |
| Tp0479 (S)<br>Tp0479 (N) | 510890..510351<br>511161..510487 | □□ □□<br>□□ □□ | ☒ ☒<br>☒ ☒ | Putative membrane<br>protein | 79.9% |
| Tp0548 (S)<br>Tp0548 (N) | 593141..594457<br>593270..594574 | ☒ □□<br>□□ □□ | ☒ ☒<br>☒ ☒ | Putative OMP; FadL<br>homolog | 94.5% |

<sup>1</sup>As annotated in NC\_021490.2 and NC\_021508.1
<sup>2</sup>SS14 Tp0006 sequence is encompassed by Tp0006 and Tp0007 ORFs in the Nichols strain. Neither Nichols' annotated Tp0006 or Tp0007 are represented in the array.
<sup>4</sup>No amplification of the target gene.
<sup>5</sup>Also annotated in the *T. pallidum* DAL-1 strain <sup>44</sup>
<sup>6</sup>Not annotated in the Nichols strain by any pipeline.

## DISCUSSION

Advancements in genomics, transcriptomics, and proteomics approaches have recently led to a significant increase in the number of *T. pallidum* genome sequences ^28^ and to the definition of a hierarchy of highly to poorly expressed genes at both the mRNA and protein level in treponemes grown *in vitro* or recovered from experimentally infected rabbits ^42,45–47^. Comparative analyses of the 514 assembled *T. pallidum* genomes sequenced to date from all continents, for example, showed that currently circulating strains fall either within the Nichols or the SS14 clades of this pathogen ^28^. Such evidence led to including SS14- and Nichols-specific targets in our PPA and the core proteome shared by both strains. Furthermore, the recent re-sequencing of both the Nichols and SS14 laboratory isolates allowed the use of updated annotations to define the boundaries of the individual ORFs cloned into the plasmid library for IVTT ^48,49^. The choice of a PPA containing the complete *T. pallidum* proteome was also supported by gene expression studies showing detectable messages for virtually all annotated genes and by the results of mass spectrometry-based studies collectively demonstrating that 90% of the *T. pallidum* annotated proteome is translated. Not all *T. pallidum* antigens are however represented in the array ^42,45–47^. Not represented targets are reported in Table S1. The TprK/Tp0897 putative OMP was purposely omitted due to its intra-strain hypervariability ^50,51^. Although a recombinant TprK fragment corresponding to the protein less variable amino-terminus (aa 29/AQV-274/ALA; carrying two predicted invariant conserved surface-exposed loops) was spotted on the arrays as a control, and this target was significantly recognized in rabbits infected with either strain (Table S1), we did not perform a formal analysis of reactivity, given that no data could be collected on the response to the missing portion of the protein. Ongoing work using a phage immunoprecipitation/sequencing approach (PhIP-Seq) employing a phage library of TprK variable and conserved sequences based on deep sequencing of the *tprK* gene will define which TprK epitopes elicited a significant humoral immunity in these infected animals.

This study attempted to close a significant knowledge gap in syphilis research by analyzing the longitudinal development of antibodies to each *T. pallidum* protein during infection and how acquired immunity changes in response to treatment. This goal was also pursued by McKevitt *et al.*^52^ who admirably produced an 882-protein array based on the Nichols genome sequenced in 1998 by expressing each single antigen in *E.coli* ^53^. Upon testing this early array with sera from long-term (86 days) Nichols-infected rabbits, however, only 106 immunoreactive antigens were identified, while our study identified 213 reactive proteins in N.LT rabbits. Aside from the lower number of targets on McKevitt’s array ^52^ compared to our PPA, this discrepancy might partially be attributable to false-negative results due - as acknowledged by the authors - to poorly expressed target proteins in a system that was still reliant on *E. coli* for protein expression rather than on IVTT. The authors also alluded to PCR-generated errors in their plasmid clones that might have altered the immunogenicity of target epitopes to explain why *T. pallidum* antigens known to be immunogenic during infection failed to be detected in their study ^52^. Regardless, a comparison between the thirty most significantly reactive antigens in both studies (^52^ and Table 1) showed very similar results, with 20 shared targets, and only five proteins (Tp0872/flagellar filament capping protein; Tp0259/LysM peptidoglycan binding protein; Tp0594/hypothetical protein; and Tp0737/periplasmic binding protein) recognized by our PPA but not by the earlier array, and six targets (Tp0453/concealed OMP; Tp0272/FlgE flagellar hook; Tp0841/DegQ endoprotease; Tp0369/BamD; Tp0468/tetratricopeptide domain protein; and Tp0433/hypothetical protein) recognized by both arrays, but associated to higher reactivity in our study. In our study, no differences were seen when signals generated by IVTT proteins and control recombinant proteins expressed in *E. coli* were seen (Fig.S4).

Of the three targets currently used in TTs based on recombinant proteins, both the 47 kDa lipoprotein/Tp0574 (TpN47) and the 17 kDa lipoprotein/Tp0435 (TpN17) were among the most highly recognized antigens by all N.LT and S.LT rabbits, also in agreement with the McKevitt study ^52^. However, the 15kDa lipoprotein/Tp0171 (TpN15) was not highly immunoreactive in infected animals in either study. Here, the N.LT and S.LT rabbit sera became significantly positive to TpN15 between day 40 and day 50 post-inoculation (Table S1), which does not fully support the use of this antigen for early diagnosis, despite being highly expressed, remarkably conserved across *T. pallidum* strains and subspecies, and sequence-specific for the agents of human treponematoses ^54^. Several clinical studies, however, support the validity of TpN15 as a serodiagnostic target ^55–63^, and this discrepancy could be due to differential immunity that develops in the rabbit compared to the natural human host. We also identified several antigens worth investigating as diagnostic candidates in addition to or in substitution of immunodominant lipoproteins to increase the sensitivity of TTs (Table 1). Among those, the hypothetical transcriptional regulator, Tp0772, elicited a fast and significantly high response in enrolled animals. Significant reactivity to this target in human sera was reported by Brinkman *et al.* ^64^, albeit using a limited number of specimens. The Tp0768/TmpA lipoprotein could be another suitable target. In a previous study, using 120 patient sera and a minimal protein array also containing Tp0768, we showed a significant correlation in reactivity between Tp0768 and both the TpN17 and TpN47 lipoproteins ^30^. However, reactivity to TmpA was generally lower than both TpN17 and TpN47 and the spread of values was greater ^30^. The possible use of Tp0768 in TTs is also supported by the results of studies published by Brinkman *et al.* ^64^ and by the van Embden group that analyzed ∼70 sera from patients at different stages of syphilis over two studies ^65,66^. The possible use of Tp0277, Tp0319, Tp0326, Tp0453, Tp0684, and Tp0768 was also supported by Runina *et al.* ^67^ and by work from other investigators as well ^68^.

The near absence of an IgM response in all sera from the N.LT, N.RI, S.LT, and S.RI animals (Fig.S3) was surprising, as we expected IgM reactivity to at least partially overlap that of the IgG-positive targets. The lack of IgM signal cannot be attributed to the lack of protein target spotted on the array or to the inability of IgM to bind the target, as positive control features in the array were reactive, but likely to the low level of IgM in the serum samples. Equally intriguing was the difference in the number of targets recognized in S.LT animals (n = 327) compared to the N.LT rabbits (n = 207) (Fig.2). Given that both groups of animals received the same inoculum size, in lack of a better explanation, this result could be related to the different growth rate of the two strains we used. Nichols’ generation time is, in fact, 41.5 hours, while SS14 needs 56.6 to divide ^69^. This apparently minor difference causes remarkable phenotypic changes in disease manifestations in infected animals. For example, in rabbits infected ID with the same inoculum size (10^6^ cells of either the Nichols or the SS14 strain), dermal lesion progression from erythema to ulceration and then healing requires virtually half the time in Nichols-infected animals. Likewise, in animals inoculated IT with 10^7^ cells/testis of Nichols treponemes, orchitis (assessed by palpation) manifests in approximately ten days, while ∼20 days are needed in rabbits infected IT with the same inoculum of SS14 cells. Whether this difference in generation time is due to the genetic diversity existing between the two strains or to the adaptation process that the Nichols strain likely experienced during its continual propagation in rabbits since 1912, one could hypothesize that faster growth translates into an accelerated immune recognition and pathogen clearance, leading to a limited array of antigens recognized before most pathogen cells are eliminated. This hypothesis could also be consistent with the work by McKevitt *et al.* ^52^ discussed above, where a treponemal inoculum 67 times higher than ours (4×10^8^ Nichols cells/rabbit) was entirely administered IT to their rabbits, potentially leading to fewer antigens being recognized in their Nichols-infected rabbits compared to ours due to a fast host response induced by an abnormally large inoculum. Although a comparison between the results of NTTs and TTs in both studies could have corroborated our hypothesis, such data were not reported by McKevitt *et al.* ^52^.

New cases of syphilis occur primarily in areas characterized by poor access to health care due to low socioeconomic status. The availability of an effective vaccine would greatly help in reducing disease incidence in adults and newborns and dependence on antibiotics to avoid infection, particularly when post-exposure prophylaxis is considered ^70–72^. Early work on syphilis immunology in the rabbit model established that rabbits infected with *T. pallidum* for at least 3 months develop immunity to symptomatic reinfection, albeit they get re-infected if inoculated again ^73,74^. Defining the humoral nature of this “chancre immunity”, might identify potential candidates to actively pursue as components of a syphilis vaccine. As *T. pallidum* clearance from infected lesions is believed to occur mainly through phagocytosis of opsonized treponemes ^75^, surface-exposed antigens of the syphilis spirochete are the most likely vaccine candidates, and several are currently pursued to induce protective immunity. Our results suggest that antibodies to the Tp0326/BamA OMP, a protein known to be associated with biogenesis of the outer membrane ^76^ might play a role in protective immunity, as also suggested in recent work by Ferguson *et. al*^77^. The comparison of the immunity generated to OMPs in both N.LT and S.LT animals, however, also suggested that there might be a core number of OMPs sufficient to induce partial protection, as relatively fewer of these putative surface-exposed antigens were recognized in N.LT rabbits compared to S.LT ones (Fig.4D). Reported immunization/challenge data for some of the antigens that appear to be dispensable seem to support this hypothesis. Rabbit immunization with Tp0126/OmpW, for example, did not induce protection following infectious challenge ^78^, and neither did immunization with the putative surface-exposed lipoprotein Tp0136 ^79^, even though recent work suggest a functional role for the immunity to this antigen in the inhibition of treponemal dissemination ^80^, consistent with its role of fibronectin-binding adhesin of *T. pallidum*. With regard to the Tp0859/FadL homolog, the presence of an homopolymeric G repeat within the ORFs ^81^ suggests that this gene might undergo translational phase variation, supporting a non-essential biological role for this putative porin transporter, likely due to the presence of four additional members in this family of paralogous proteins in the *T. pallidum* proteome.

The rabbit re-infection experiment helped address a limitation of TTs, namely the continued reactivity of modern conventional TTs after successful treatment that makes them unsuitable for monitoring treatment response, relapse, or reinfection in previously treated subjects. In this context, the identification of novel *T. pallidum* antigens, whose antibodies are rapidly eliminated from the host circulation, could lead to tests that could complement the NTTs, which would be helpful in the case of serological non-responders, who do not achieve the 4-fold dilution in NTT titer necessary to declare serological cure, or serofast patients who do not serorevert even if the proper reduction in NTT titer was achieved ^15^. In our study, although reactivity to several antigens underwent a reduction over time, we could not find a specific target recognized by most infected animals that exhibited a significant reduction in reactivity between day 30 post-infection, at the time of treatment, and day 90, right before re-infection. This could also be due to the relatively short time elapsed between treatment of these animals and re-infection, and the outcome of this experiment cannot exclude the possibility that treponemal antigens could have a role in assessing successful treatment. In a previous study by Haynes *et al*. ^30^ using patient samples and a minimal proteomic array carrying only 16 antigens, we identified an association with RPR titer drop post-treatment and a reduction in the signal to Tp0679/TmpB, as previously reported in a similar study by Shouls *et al.* ^66^. Although the result reported here using experimental sera did not confirm the earlier report, our ongoing analysis of clinical samples would shed further light on the possible use of TmpB or other antigens for the monitoring of treatment efficacy.

In summary, we have adapted a robust high throughput proteome-synthesis process to two strains of the syphilis agent and have provided proof of concept that the resulting proteomic array can define antigen-specific antibody profiles from infected laboratory subjects. The application of this technology, both at the pre-clinical and clinical level, will contribute to improving our understanding of syphilis pathogenesis and immunology, and to accelerating the discovery of potential vaccine candidates and antigens for diagnostics applications. Ongoing studies in the laboratory are evaluating the humoral response to *T. pallidum* antigens using 217 sera longitudinally collected pre- and post-treatment from 120 patients with syphilis enrolled between 2019 and 2021 in five sexual health clinics in Peru. Such data will allow us to further investigate whether treponemal antigens could be used to perform disease staging and monitor response to treatment in patients with different HIV status history of infection. For our ongoing syphilis in pregnancy study (SIPS), samples from two additional cohorts of pregnant subjects with syphilis will be tested to assess whether a correlation exists between immunity to specific *T. pallidum* antigens and pregnancy outcomes.

## CONTRIBUTORS

J.J.C. and L.G. conceived of the study. L.G., E.R. A.P. and A.M.H. performed the animal work. A.O., J.V.P. and A.A.T. designed the array. J.V.P. established the clone library. C.H. and A.D.S. probed the array. J.J.C. and L.G. analyzed the data. All authors contributed to manuscript preparation and critically reviewed and approved the manuscript before submission.

## DATA SHARING STATEMENT

All reactivity data used to generate figures are reported in File S1. All datasets generated by probing each array and analyzed during the current study are available upon request to both Dr. Campo and Dr. Giacani.

## ACKNOWLEDGEMENTS

The authors are grateful to Angela Yee (Antigen Discovery Inc.) for critical review of the manuscript. The funders of the study had no role in the study design, data collection, data analysis, data interpretation, writing of the report, or in the decision to submit the paper for publication.

---

**Figure S1.**
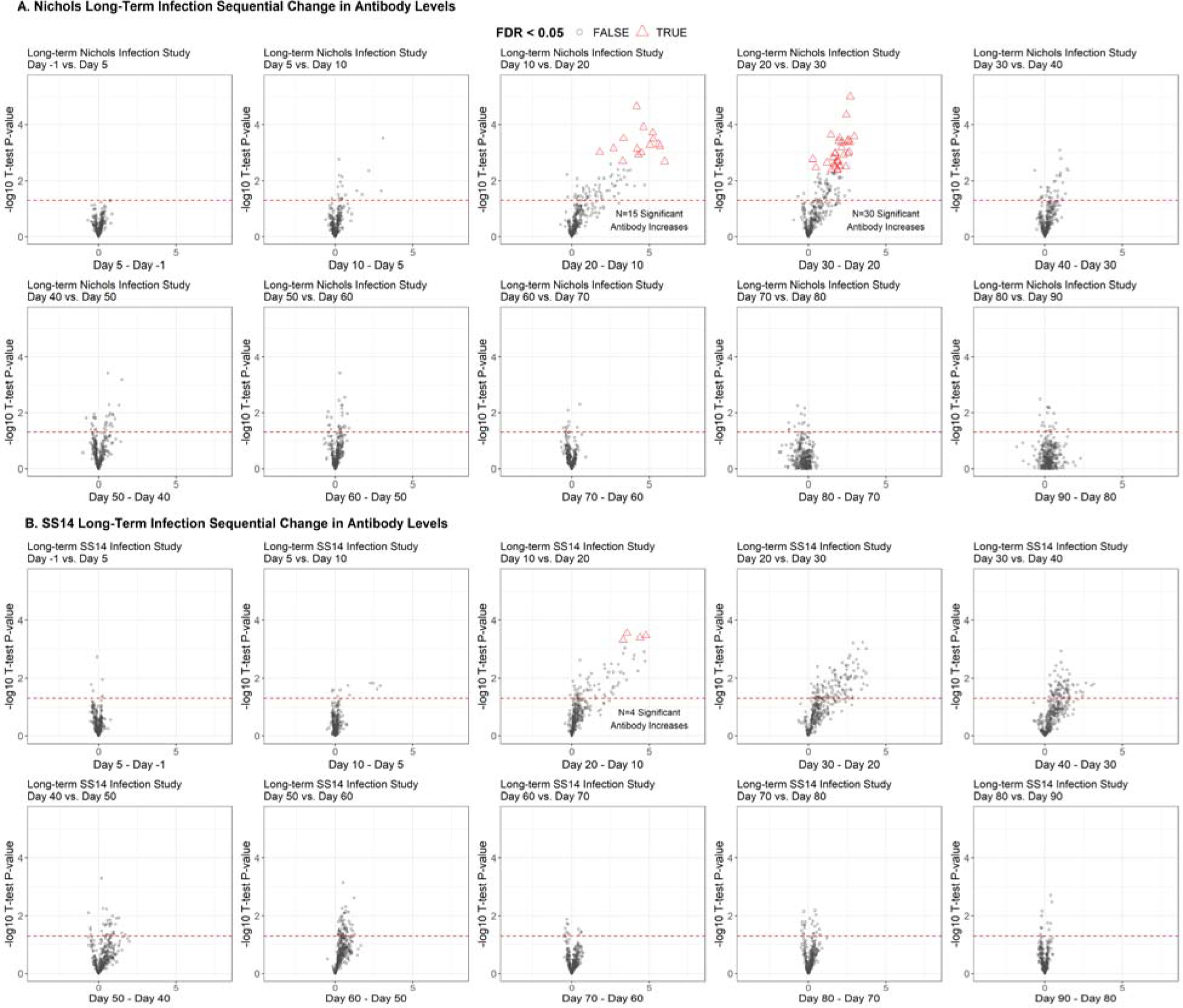
Longitudinal trajectory of IgG binding levels between sequential time points in long-term (N.LT and S.LT) infected rabbits. Volcano plots showing the difference in normalized IgG binding between timepoints for each reactive protein on the array (x-axis) by the inverse log_10_ P-value of paired Student’s T-tests. In each plot, the horizontal red dashed line represents an unadjusted P-value of 0.05, and all values above the red line are below 0.05. After adjustment for the false discovery rate, antibody responses that remained statistically significant were highlighted as red triangles. **(A)** Sequential time point differences (Day −1 vs day 5; day 5 vs day 10, day 10 vs day 20 and so on) for long-term Nichols-infected animals. **(B)** Sequential time point differences for long-term SS14-infected animals.

**Figure S2.**
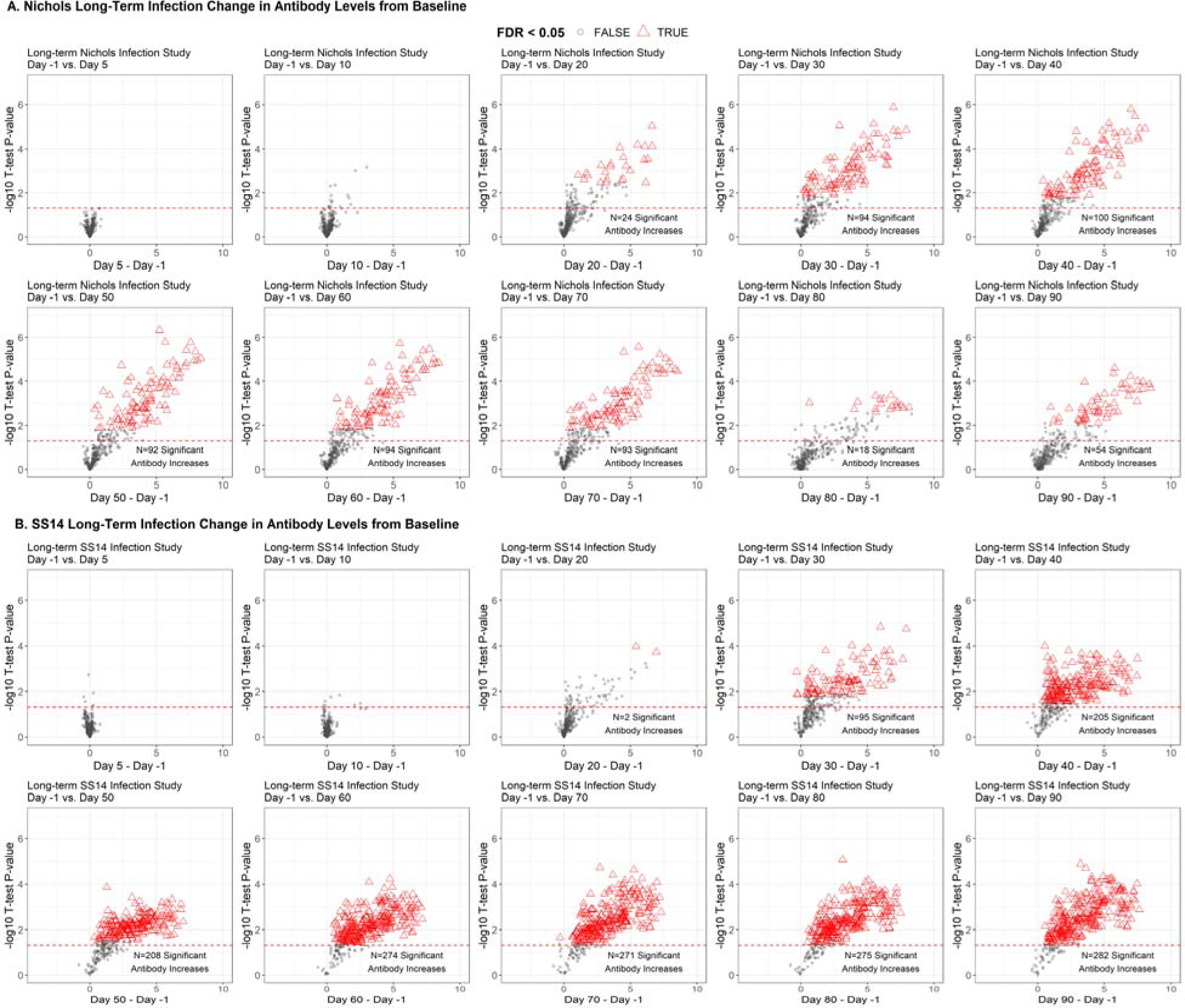
Longitudinal trajectory of IgG binding levels versus baseline in long-term (N.LT and S.LT) infected rabbits. Volcano plots showing the difference in normalized IgG binding of each post-infection time point versus the pre-infection (Day −1) time point for each reactive protein on the array (x-axis) by the inverse log_10_ P-value of paired Student’s T-tests. In each plot, the horizontal red dashed line represents an unadjusted P-value of 0.05, and all values above the red line are below 0.05. After adjustment for the false discovery rate, antibody responses that remained statistically significant were highlighted as red triangles. **(A)** Sequential time point differences for long-term Nichols-infected animals vs day −1. **(B)** Sequential time point differences for long-term SS14-infected animals vs day −1.

**Figure S3.**
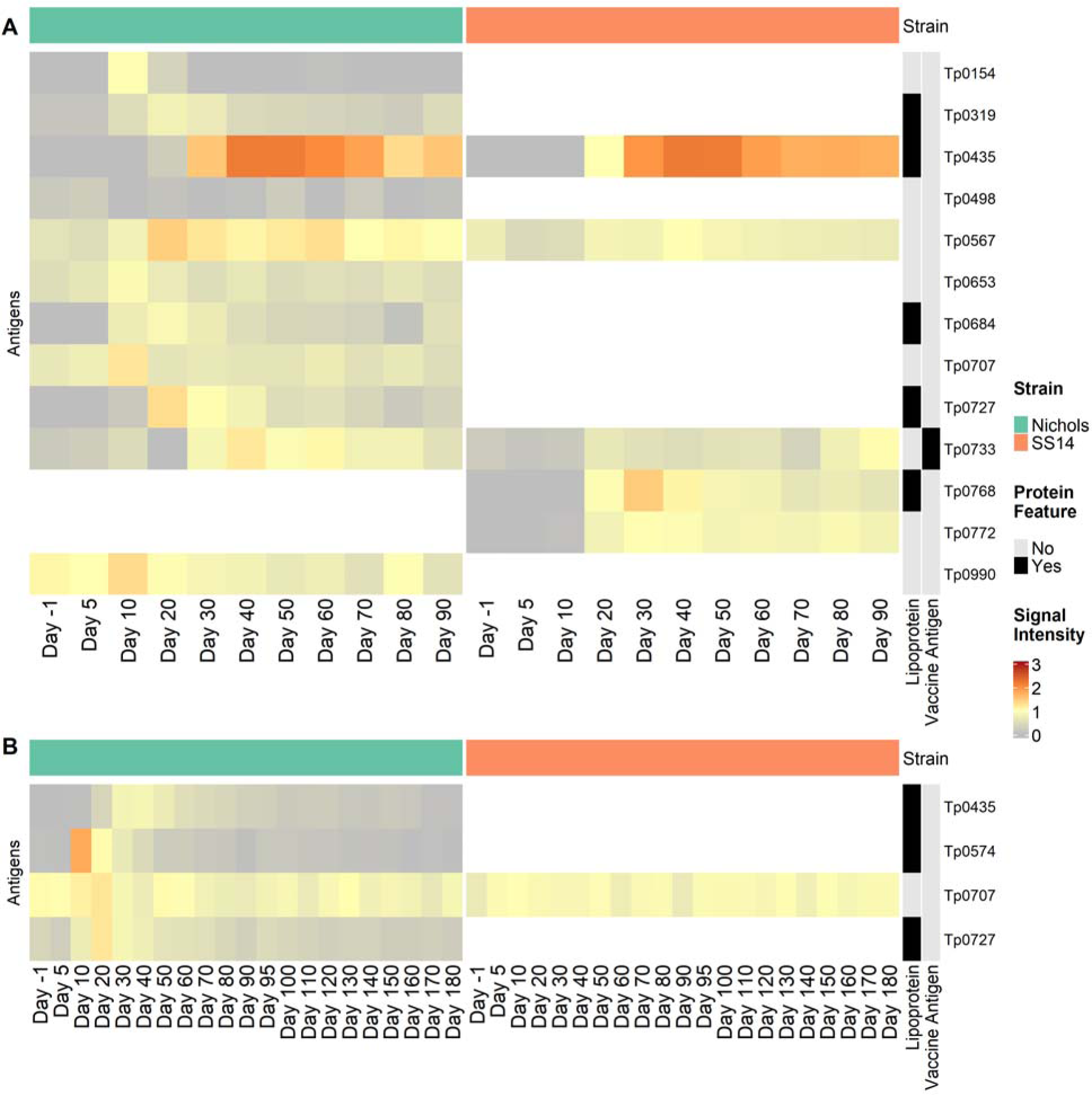
IgM antibody binding profile in LT and RI rabbit groups. **(A)** Heatmap of the 13 total antigens overall recognized in long-term Nichols-infected animals (left side, green header), and SS14-infected rabbits (right side, orange header) over the 90-day infection period. Only the *T. pallidum* proteins with significant immune reactions, or at least 4 out of 6 rabbits that responded to the protein in either strain group, were included in the heatmap. The category of each protein as a lipoprotein or possible vaccine candidate is shown in black and grey at the right of the heatmap**. (B)** Heatmap of the four antigens overall recognized in Nichols-infected animals (left side, green header), and SS14-infected rabbits (right side, orange header) over the 180-day infection/treatment/re-infection experiment. Treatment occurred at day 30 post-inoculation, and reinfection with the homologous strain occurred 60 days post-treatment. Only the *T. pallidum* proteins with significant immune reactions, or at least four out of six rabbits that responded to the protein in either strain group, were included in the heatmap. The category of each protein (lipoprotein or a vaccine candidate) is shown in black and grey at the right of the heatmap.

**Figure S4.**
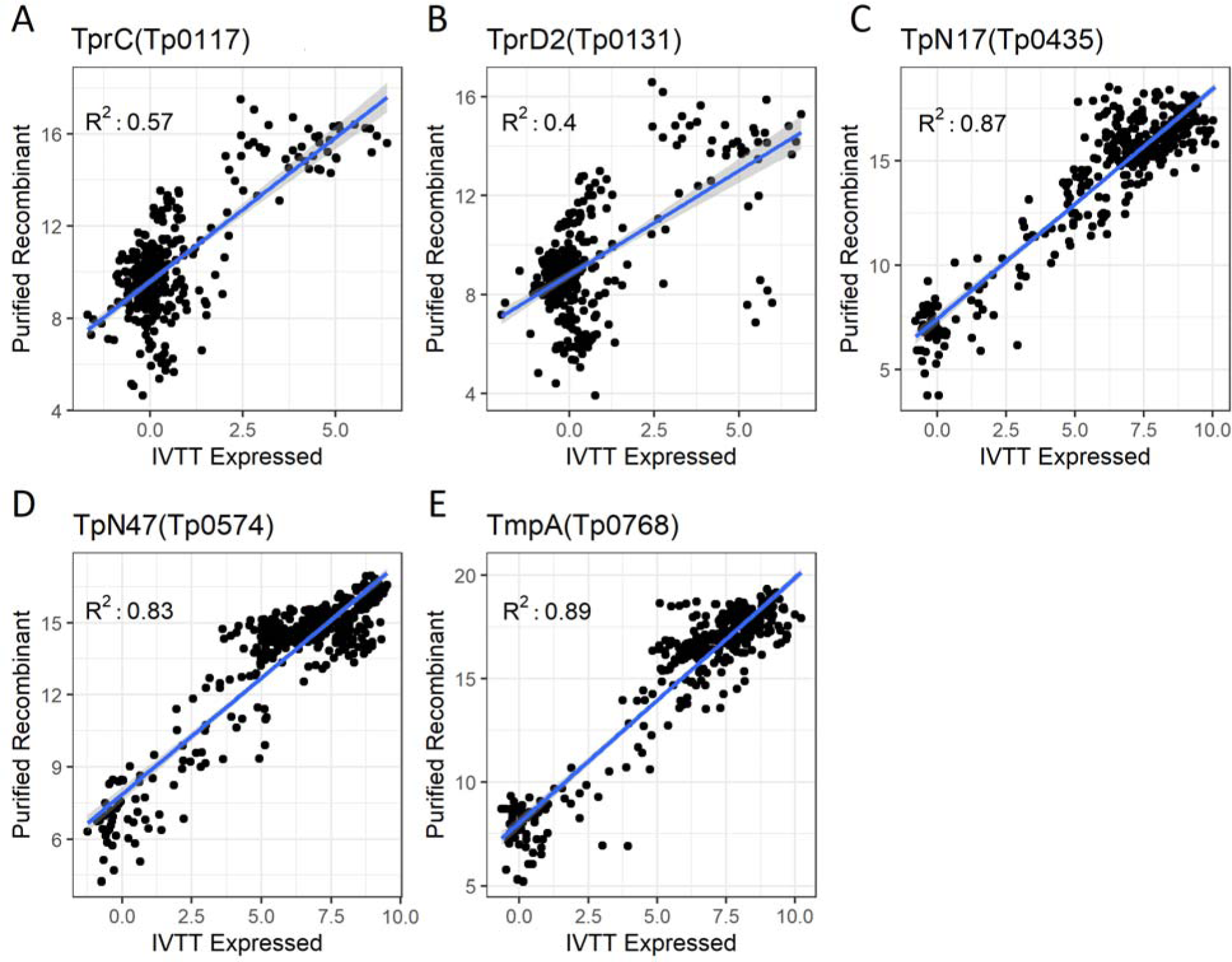
Correlation between serum reactivity of IVTT expressed protein and control recombinant proteins expressed in *E. coli*. **(A-E)** Pearson correlations between signal to IVTT-expressed and purified recombinant protein TprC (Tp0117), the SS14-specific TprD2 (Tp0131) antigen, TpN17 (Tp0435), TpN47 (Tp0574), and TmpA (Tp0768). R values are reported within individual panels.

